# Role of Water-bridged Interactions in Metal Ion Coupled Protein Allostery

**DOI:** 10.1101/639468

**Authors:** Xingyue Guan, Cheng Tan, Wenfei Li, Wei Wang, D. Thirumalai

**Author notes:** **For correspondence:** (WW); (DT). These authors contributed equally to this work (Xingyue Guan, Cheng Tan).

## Abstract

Allosteric communication between distant parts of proteins controls many cellular functions, in which metal ions are widely utilized as effectors to trigger the allosteric cascade. Due to the involvement of strong coordination interactions, the energy landscape dictating the metal ion binding is intrinsically rugged. How metal ions achieve fast binding by overcoming the landscape ruggedness and thereby efficiently mediate protein allostery is elusive. By performing molecular dynamics simulations for the Ca^2+^ binding mediated allostery of the calmodulin (CaM) domains, each containing two Ca^2+^ binding helix-loop-helix motifs (EF-hands), we revealed the key role of water-bridged interactions in Ca^2+^ binding and protein allostery. The bridging water molecules between Ca^2+^ and binding residue reduces the ruggedness of ligand exchange landscape by acting as a lubricant, facilitating the Ca^2+^ coupled protein allostery.

Calcium-induced rotation of the helices in the EF-hands, with the hydrophobic core serving as the pivot, leads to exposure of hydrophobic sites for target binding. Intriguingly, despite being structurally similar, the response of the two symmetrically arranged EF-hands upon Ca^2+^ binding is asymmetric. Breakage of symmetry is needed for efficient allosteric communication between the EF-hands. The key roles that water molecules play in driving allosteric transitions are likely to be general in other metal ion mediated protein allostery.

## Introduction

Calmodulin (CaM), a versatile calcium (Ca^2+^) sensing protein expressed in all eukaryotic cells, is involved in a bewildering range of intracellular signaling processes ***Clapham (2007***). Examples include activation of kinases ***Osawa et al. (1999***); ***Wang et al. (2019***), muscle contraction ***Walsh (1994***), gene regulation ***Means (1994***), signal transduction ***Gunther et al. (1993***); ***Liu et al. (2017***); ***Zhao et al. (2019***), and apoptosis ***Jiang et al. (1996***). Binding of Ca^2+^, whose intracellular and extracellular concentrations differ by several orders of magnitude, to CaM results in a large conformational change, leading to the exposure of hydrophobic residues that serve as recognition sites for target proteins. CaM is composed of two nearly symmetric globular domains connected by a flexible central helix. Each domain consists of two helix-loop-helix motifs, termed EF-hands (Fig. 1A,B), which are found in a large number of calcium-sensing proteins ***Gifford et al. (2007***); ***Lewit-Bentley and Réty (2000***). The EF-hands chelate Ca^2+^, resulting in coordination to seven ligands arranged in a pentagonal bipyramid geometry (often involving one water molecule and five residues) with the negatively charged residues in the loop ***Lewit-Bentley and Réty (2000***) (Fig.1B). In the apo state of CaM (Fig. 1A), the helices in the EF-hand motif are arranged in an anti-parallel manner. Upon binding of Ca^2+^, the helices undergo substantial rearrangement into a nearly perpendicular conformation, exposing the hydrophobic sites which enable recognition and subsequent activation of target proteins ***Gifford et al. (2007***); ***Grabarek (2006***); ***Nelson and Chazin (1998***). Through such a mechanism, CaM initiates a variety of cellular processes.

**Figure 1.**
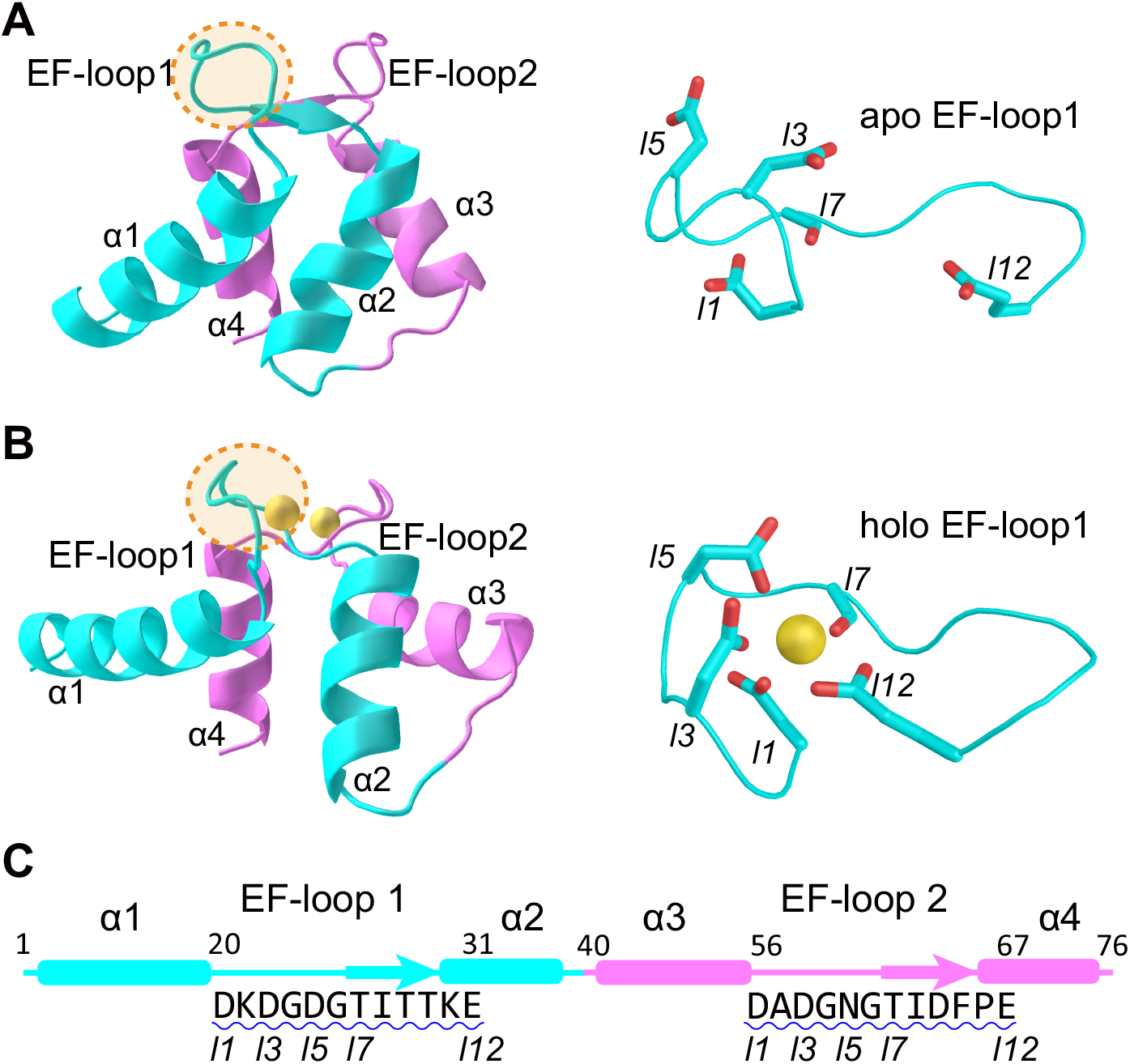
Structure and sequence of nCaM and the Ca^2+^ binding EF-loops. (A, B) Structure of nCaM in the holo state (A) and apo state (B). Helix-loop-helix motifs *α*1–EF-loop1–*α*2 (cyan) and *α*3–EF-loop2–*α*4 (violet) comprise two EF-hands in the N-terminal domain of Calmodulin. Yellow spheres correspond to Ca^2+^ ions. The details for one of the allosteric site in the loop region were also shown (right). (C) The sequence of nCaM and corresponding secondary structures. Ca^2+^ binding loops are underlined with blue wavy lines. The native ligands are marked with “*ln*” to indicate their position in the EF-loop.

In a more general context, Ca^2+^-triggered conformational changes in CaM is an example of allostery, which describes the structural changes that occur in enzymes and molecular machines at distances far from the site at which a ligand binds ***Tsai et al. (1999***); ***Thirumalai et al. (2019***); ***Saavedra et al. (2018***). Nature utilizes such functional motions, which are triggered by binding of cofactors to specific sites in proteins and RNA, to execute a variety of functions ***Hyeon et al. (2006***); ***Weinkam et al. (2012***); ***Li et al. (2011***); ***Ferreiro et al. (2011***); ***Motlagh et al. (2014***); ***Mugnai et al. (2020***); ***Li et al. (2019***); ***Wodak et al. (2019***); ***Wang et al. (2018***); ***Raguimova et al. (2020***); ***Li et al. (2014***). Binding of cofactors at a certain site (allosteric site) induces structural rearrangements at a distant site (regulated site) in the enzyme, thereby modulating the downstream activity. By such allosteric (action at a distance) movements, small local changes are amplified over long distances, which allows for control and regulation of cellular signaling and functions ***Csermely et al. (2010***). Divalent metal ions, e.g., Ca^2+^ and Zn^2+^, have been widely utilized as effectors to trigger the allosteric cascade. However, due to the involvement of strong coordination interactions, the energy landscape dictating the metal ion binding and unbinding is intrinsically rugged. Despite the biological importance of metal ion coupled protein allostery, two key questions remain elusive, including: i) What strategy do metal ions utilize to overcome the ruggedness of the binding landscape and achieve rapid binding/unbinding? and ii) What physical interactions enable the propagations of the metal ion triggered allosteric signal to regulate the downstream processes? Because of the availability of rich experimental data and the typical allosteric features, CaM is an ideal system for answering these questions computationally ***Li et al. (2014***); ***Linse et al. (1991***); ***Barton et al. (2002***); ***Junker et al. (2009***); ***Chen and Hummer (2007***); ***Price et al. (2011***); ***Tripathi and Portman (2009***); ***Itoh and Sasai (2011***); ***Ovchinnikov and Karplus (2012***); ***Li et al. (2015***); ***Shukla et al. (2016***); ***Tamura and Hayashi (2015***); ***Chen and Hummer (2007***); ***Park et al. (2008***); ***Malmendal et al. (1999***); ***Evenäs et al. (2001***); ***Ohashi et al. (2011***); ***Xiong et al. (2010***); ***Zhang et al. (2008***); ***Kukic et al. (2016***); ***Grabarek (2005***).

Here, we used the bias-exchange metadynamics method ***Piana and Laio (2007***) to investigate the Ca^2+^ binding coupled conformational changes in CaM domains by explicitly modeling Ca^2+^ in the simulations. We extracted the free energy landscape of the Ca^2+^ binding coupled conformational changes of CaM domains at atomic level and demonstrated the full picture of the allosteric motions of CaM ***Fiorin et al. (2005***); ***Chen and Hummer (2007***); ***Likić et al. (2003***); ***Zhang et al. (2007***). Our results revealed the key role of water-bridged interactions during Ca^2+^ binding and protein allostery. The bridging water molecules between Ca^2+^ and binding residues reduce the free energy barrier of ligand exchange landscape, and therefore the landscape ruggedness, by acting as a lubricant. This enables efficient Ca^2+^-ligand coordination and allosteric motions in the CaM. We propose that the water-bridged coordination is a general mechanism utilized in metal-coupled folding and allosteric communication in proteins. We also showed that Ca^2+^ binding leads to rotation of the EF-hand helices with the hydrophobic core as the pivot, a structural change that should precede recognition by target proteins for signal transduction. In addition, The molecular simulations revealed obvious asymmetry in the allosteric coupling of the symmetrically paired EF-hands: the EF-hand 1 (EF_1_) binds Ca^2+^ in a sequential manner by chelating to the N-terminal residues followed by coordination with the central residues, and finally to the C-terminal residues. In contrast, chelation of Ca^2+^ to the EF-hand 2 (EF_2_) is initiated either by interactions with the residues in the N- and/or C-terminal residues followed by coordination to the central residues. Similar results were obtained for the cCaM. Such a symmetry breaking process is likely involved in larger multi-domain complexes, such as the bacterial chaperonin, GroEL, in which ligand binding induces asymmetric response in different subunits.

## Results and discussions

### Ca^2+^ binding is coupled to conformational changes of the EF-hand motifs

We use well-tempered bias-exchange metadynamics ***Piana and Laio (2007***); ***Barducci et al. (2008***) and the corresponding reweighting techniques ***Bonomi et al. (2009***) to extract the free energy landscape projected onto physically motivated multi-dimensional reaction coordinates characterizing Ca^2+^ binding and the conformational change. Quantitative analysis of the changes in these co-ordinates allows us to infer the mechanism of coupling between Ca^2+^ binding, the role of water, and the allosteric conformational changes in calmodulin. In particular, we use the “path collective variables” ***S***_*α*_ (*α* = 1, 2, 3, and 4), defined in the **Methods and Materials** section, to describe the conformational changes of the two EF-hand motifs. Small and large values of ***S***_*α*_ correspond to open and closed states, respectively. In order to assess the consequences of Ca^2+^ binding, we use the native coordination numbers ***N***_*α*_ (**Methods and Materials**), representing the number of native ligands (oxygen atoms of residues that are coordinated with Ca^2+^ in the native holo structure) that bind to Ca^2+^ during the allosteric transitions for the two EF-hand motifs, respectively. Figure 2 shows the free energy landscapes, *F* (***N***_1_, ***S***_1_) and *F* (***N***_2_, ***S***_2_), of the nCaM. For both EF_1_ and EF_2_, the conformational change of nCaM is tightly coupled to the extent of Ca^2+^ binding as indicated by the conformational distributions at different ***N***_1_ and ***N***_2_ values. When ***S***_*α*_ (*α* = 1 or 2) is large and ***N***_*α*_ (*α* = 1 or 2) is small, the closed state is more stable. Whereas when the native ligands are fully coordinated to Ca^2+^ the open structure is more stable (Fig. 2 and Fig. S1). Interestingly, even with full coordination of native ligands (***N***_1_ = 6) to Ca^2+^, the EF_1_ samples a wide range of conformations as assessed by ***S***_1_ fluctuations, suggesting the conformational plasticity of the open state. Conformational plasticity for the Ca^2+^ bound state, which was also observed in experiments, could be the major reason that the EF-hand motifs recognize and bind a variety of target proteins ***Chou et al. (2001***); ***Vigil et al. (2001***).

**Figure 2.**
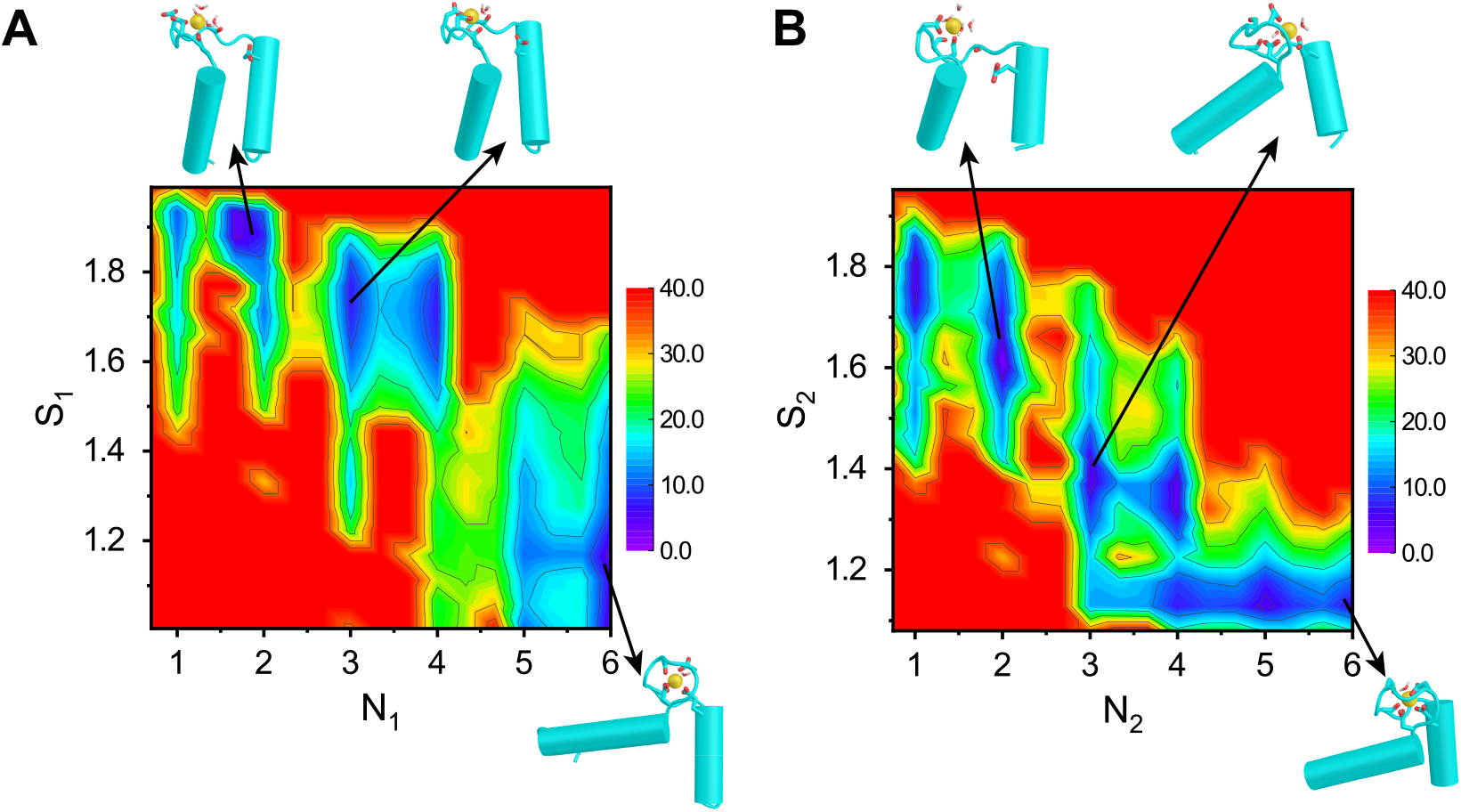
Free energy profiles of the Ca^2+^ coupled conformational transitions of EF_1_ (A) and EF_2_ (B) projected onto the collective variables (***N***_1_, ***S***_1_) and (***N***_2_, ***S***_2_), respectively. Representative conformations of the major basins in the landscape are also shown. The unit of the free energy is kJ/mol.

Although the two EF-hand motifs have similar structures, comparison of Fig. 2A and Fig. 2B shows that there are noticeable differences in the free energy landscapes upon Ca^2+^ binding. For example, when three or four ligands bind to Ca^2+^, the conformation of EF_2_ is poised to make a transition to the open-like structure, whereas EF_1_ remains in the closed-like structure. Detailed analysis shows that during the conformational transition, the two EF-hand motifs are tightly coupled. The open conformation of EF_2_ is stabilized when EF_1_ is in the open conformation with most of the associated native ligands bound to Ca^2+^. Thus, the behavior of EF_2_ is affected by the conformation of EF_1_ through long range allosteric interactions or action at a distance between the two EF-hands, which will be discussed further in later subsection. The observed differences in the allosteric communication are reasonable since the two EF-hand motifs have different amino-acid sequences in the Ca^2+^ binding loop. Consequently, the ensembles of structures with three or four native ligands bound to the Ca^2+^ should exhibit substantial differences in the two EF-motifs. Such an asymmetry in the Ca^2+^ binding and conformational transitions of the symmetrically arranged EF-hand pair in CaM domains likely plays an important role in signaling.

### Step-wise dehydration of Ca^2+^ and ligand coordination are cooperatively coupled

Our metadynamics simulations allow us to construct the changes in the coordination states of Ca^2+^, including the role that water and potential non-native ligands (those that are not present in the native open structure) play during the closed → open transition. Figure 3 shows the average of the total coordination number, ***N***_***T***_ (sum of the number of water molecules and the number of ligands bound to Ca^2+^), as a function of ***N***_*α*_ (*α* = 1, 2) for the nCaM. At a low value of ***N***_1_ (Fig. 3A), approximately five water molecules are coordinated to Ca^2+^. As ***N***_1_ increases, water molecules are expelled, showing that dehydration occurs in steps. The number of water molecules expelled from Ca^2+^ and replaced by native ligands depends on ***N***_1_. Upon binding of the first native ligand (***N***_1_ = 1), Ca^2+^ loses approximately two water molecules (Fig. 3A). Notably, the first ligand is often the aspartate, which has two carboxyl oxygens. Although only one carboxyl oxygen atom is coordinated to the Ca^2+^ in the final native coordination structure, the second carboxyl oxygen atom may also bind to the Ca^2+^ in the early stage of the coordination (Fig. 3A, gray). In comparison, one water molecule is ejected when ***N***_1_=2, 3, and 4. With the binding of the last two native ligands, the number of coordinated water molecules remains unchanged. In the holo state (***N***_1_ = 6), only one water molecule is bound to the Ca^2+^, which makes the coordination number saturate at ***N***_***T***_ = 7. (Fig. 3A).

**Figure 3.**
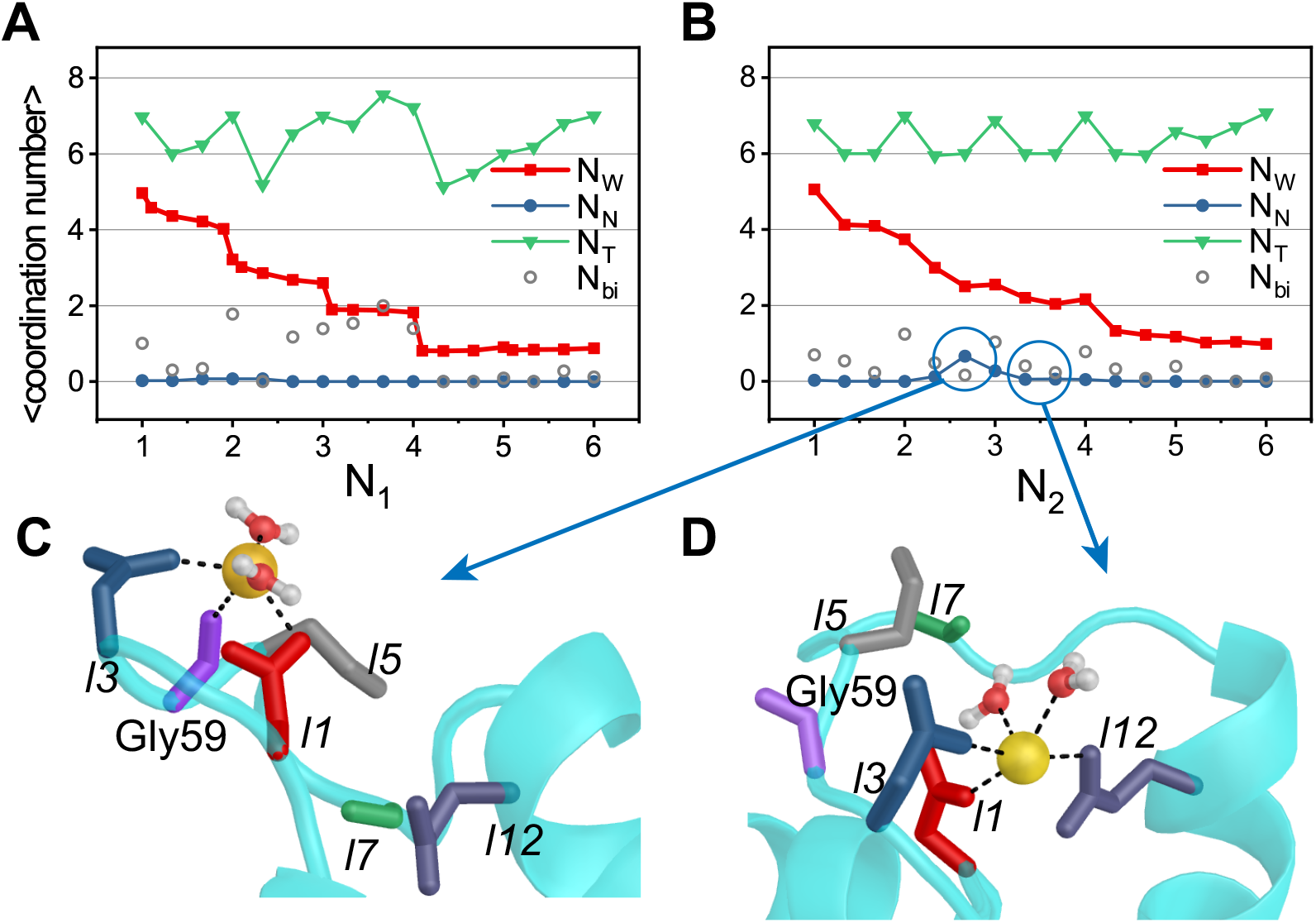
Number of coordinated water molecules (***N***_*W*_), non-native ligands (***N***_***N***_) as well as the total coordination number (***N***_***T***_) as functions of ***N***_1_ and ***N***_2_ for EF_1_ (A) and EF_2_ (B). Only one carboxyl oxygen atom of aspartate is coordinated to the Ca^2+^ in the native coordination structure. Nevertheless, the second carboxyl oxygen atom may also bind to the Ca^2+^ in the intermediate state of the coordination. The average number of such non-native bidentate ligands is also shown in panels A and B (N_*bi*_, gray). (C) The non-native ligand Gly59 (purple) coordinated to Ca^2+^ when ***N***_2_ = 2 ∼ 3. (D) When ***N***_2_ = 3, Glu67 (*l*12, purple) is coordinated to Ca^2+^, and the bond formed by Gly59 and Ca^2+^ is broken.

A similar process, with one crucial difference, occurs as Ca^2+^ coordination drives the conformational changes in EF_2_. During the binding of the native ligands (***N***_2_ ≤ 3 in Fig. 3B), Ca^2+^ becomes transiently coordinated to non-native ligands, notably the side-chain oxygen atoms of Gly59. The carboxylic oxygen of Gly59 has a probability of ∼ 0.67 to come into the first ligand shell of Ca^2+^ during the binding of the third native ligand (see Table 1 and Fig. 3C; the probability of the coordination is defined in the Supplementary Information). As the binding process and conformational changes proceed further, the non-native ligands (oxygen atoms from Gly59) are replaced by the native ligands. Figures 3C,D explicitly show that non-native coordination of Gly59–Ca^2+^ is replaced by Glu67–Ca^2+^, a ligand present in the native holo state. Such mis-ligation and ligand-exchange may be necessary to minimize the free energy cost in the sharp dehydration of metal ion during the binding process ***Li et al. (2008***); ***Lee and Lim (2011***); ***Wilson et al. (2004***).

**Table 1.** Probabilities of non-native ligand coordination and water-bridged native ligand coordination. O_*w*_ refers to oxygen in water. ***N***_*α*_ is the number of coordinated native ligands.

| $\text{EF}_1$ | $N_1=3$ | $N_1=4$ | $N_1=5$ |
| --- | --- | --- | --- |
| $\text{Ca}^{2+}\text{-O}_w\text{-Thr26}$ | 0.86 | 0.70 | 0.08 |
| $\text{Ca}^{2+}\text{-O}_w\text{-Glu31}$ | 0.67 | 0.78 | 0.14 |
| $\text{EF}_2$ | $N_2=3$ | $N_2=4$ | $N_2=5$ |
| $\text{Ca}^{2+}\text{-O}_w\text{-Asn60/Thr62}$ | 0.10 | 0.31 | 0.65 |
| $\text{Ca}^{2+}\text{-Gly59}$ | 0.67 | 0.05 | 0.02 |

### Asymmetry in the Ca^2+^ coordination pathways of the EF-hand pair

The results presented above strongly suggest that the conformational changes in the nCaM are coupled to the binding of Ca^2+^ with water playing a crucial role. Therefore, it is of interest to further clarify the order of binding of residues that are coordinated to Ca^2+^ in detail. Due to the nature of the metadynamics simulations, the kinetic information can only be indirectly gleaned using the present simulations. In particular, we can approximately extract the binding order based on the correlation analysis, as done in previous study ***Li et al. (2008***).

Figure 4A shows the coordination probability, 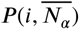, (defined in Supplementary Information) for each ligand bound to the Ca^2+^ ion in the native state with the total native coordination number being one to six for the EF_1_ of nCaM. Using 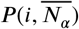 we can extract the binding order of the native ligands of EF_1_ to Ca^2+^. The representative structures are also shown (Fig. 4B). Similar results for EF_2_ are shown in Fig. 4C,D. For EF_1_, Ca^2+^ tends to bind first to the three negatively charged residues Asp20, Asp22, and Asp24 at the N-terminal part of the EF-loop. This observation is not surprising since negatively charged states of these residues contribute to the preferred binding and localization of Ca^2+^. With the progression of the Ca^2+^ chelation, several coordination bonds between the Ca^2+^ and water molecules need to be ruptured (Fig. S2), which may lead to the high free energy barrier between different coordination stages as shown in the two-dimensional free energy profiles (Fig. 2).

**Figure 4.**
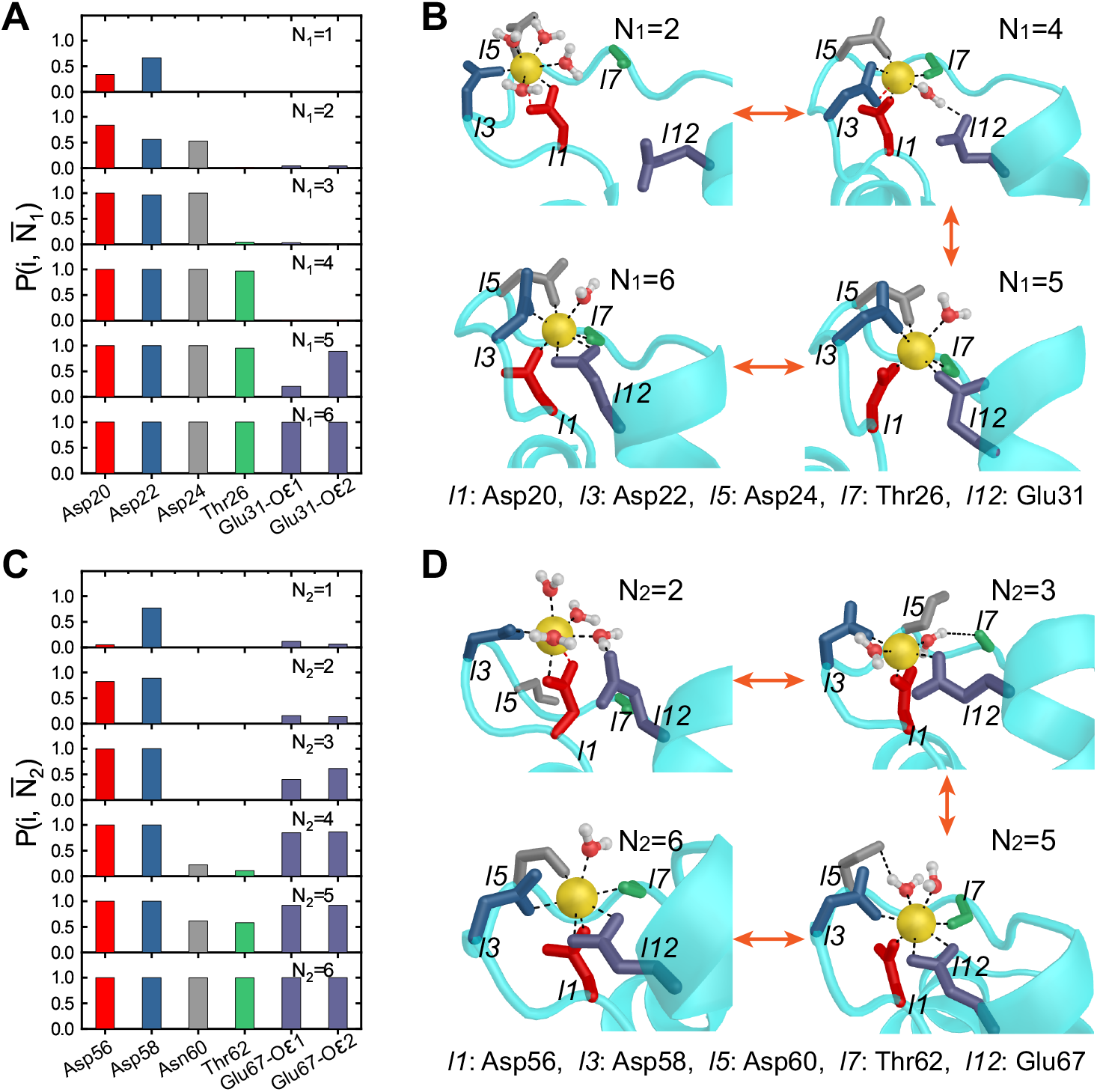
The Ca^2+^ binding sequence of EF_1_ and EF_2_. (A) Coordination probability 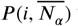 for the six native amino acids residues in EF_1_. The residues coordinated to Ca^2+^ are shown in different colors. (B) Schematic representations of the binding process of the native ligands. (C) and (D) show the binding sequence of the EF_2_.

Another possible reason for such a high free energy barrier is the dehydration of Ca^2+^ involving rupture of several coordination bonds between the Ca^2+^ and water molecules ***Pavlov et al. (1998***); ***Jalilehvand et al. (2001***). The fourth native ligand, coordinated to Ca^2+^ in EF_1_, is the backbone oxygen of Thr26, which is at the central part of the EF-loop. Binding of Ca^2+^ to Thr26 results in the formation of a water-bridged interaction between Glu31 and Ca^2+^ (Fig. 4B). In this structure, the oxygen of the water molecule is coordinated to Ca^2+^ as a ligand. At the same time, it forms a hydrogen bond with the side-chain oxygen from Glu31. We note that such a water-bridged coordination structure is quite similar to the intermediate structure identified in the crystal structure of a CaM mutant ***Grabarek (2005***). When the bridging water is finally expelled, the last two native ligands, side-chain oxygen atoms from Glu31, are coordinated to Ca^2+^. Overall, our results indicate a step wise binding of Ca^2+^ to EF_1_: first, Ca^2+^ is captured by an acidic residue (most probably Asp22); subsequently, the N-terminal part of the EF-loop plays a role in further coordination to Ca^2+^, establishing links between the incoming helix and the central *β* strands; finally, Thr26 and Glu31 are sequentially captured and bound to the Ca^2+^, leading to the native structure.

Interestingly, the results in Fig. 2 and Fig. 4 show that once one of the two oxygen atoms from Glu31 are coordinated to Ca^2+^, the most stable conformation of EF_1_ switches to the open form, suggesting the crucial role of the highly conserved Glutamic acid at the 12^*th*^ position in the Ca^2+^ binding coupled conformational transition of nCaM ***Grabarek (2006***, 2005); ***Evenäs et al. (1998***). A recent work based on nuclear magnetic resonance measurements also suggested the important role of the bidentate ligand Glu140 on the Ca^2+^ coupled conformational motions of the cCaM ***Kukic et al. (2016***).

The molecular process of Ca^2+^ binding to EF_2_ is dramatically different (Fig. 4C,D). Instead of binding to the native ligands by the sequence of N-terminal residues → central residues → C-terminal residues as in EF_1_, the Ca^2+^ binds initially to the N-terminal residues Asp56, Asp58 (or C-terminal residues Glu67), which is followed by the binding of the C-terminal residues Glu67 (or N-terminal residues Asp56, Asp58). Only at the final stage, it is bound to the central residues Asn60 and Thr62. Therefore, the Ca^2+^ binding to the EF_2_ follows the sequence N-terminal and/or C-terminal residues → central residues. The differing mechanism of coordination to Ca^2+^ probably arises from the divergence of their amino-acid sequences. For example, the “central” residues (especially at the 5^*th*^ position) in EF_1_ are more negatively charged compared to those in EF_2_. Consequently, coordination of these residues to Ca^2+^ occurs at a relatively early stage.

We also found that before Ca^2+^ binds to the central residues Asn60 and Thr62, a non-native ligand Gly59 is coordinated with Ca^2+^ (Fig. 3B,C and D). Consequently, the later steps of the Ca^2+^ binding to EF_2_ must involve ligand exchange between the non-native ligands and the native ligands. Interestingly, during the ligand exchange, water molecules bridge the Ca^2+^ and the native ligands. As we show below, water mediated coordination reduces the free energy barrier of the formation of native coordination bonds. For EF_2_, the water molecules were found during the formation of coordination of Ca^2+^–Asn60 and Ca^2+^–Thr62, which is consistent with the observation that there is a tendency to capture an extra water molecule as a ligand leading to a decreased contribution for the backbone oxygen of Thr62 ***Likić et al. (2003***). In experiments ***Grabarek (2005***), Asp64 was mutated to Asn64 for the EF_2_. Interestingly, it was shown that the mutated EF_2_ has similar binding mechanism as EF_1_, namely, the N-terminal part of the loop are coordinated to Ca^2+^ earlier than the C-terminal ligands ***Grabarek (2005***). These results suggest that the charged state of the residues to which Ca^2+^ is coordinated plays a crucial role in the mechanism of Ca^2+^ binding, and hence allostery ***Csermely et al. (2010***).

The cCaM exhibits a similar Ca^2+^ binding order except that the order of events in the EF_3_ and EF_4_ in the cCaM corresponds to those of the EF_2_ and EF_1_ in the nCaM (Fig.S3), as expected from the similarity of the sequences of the respective Ca^2+^ loops. As shown in Fig. 1 and Fig. S3, the residue at the *l*5 position of the EF_1_ (EF_4_) in the nCaM (cCaM) is charged, whereas those in the EF_2_ (EF_3_) in the nCaM (cCaM) are neutral. The stepwise dehydration of the Ca^2+^ in the cCaM was also observed (Fig. S4).

### Ca^2+^ binding-induced rotation of EF-hand helices

Structures of nCaM show that there are several hydrophobic residues located around Ile27 and Ile63 whose non-polar side-chains stack against each other to form the center of the hydrophobic core ***Chou et al. (2001***); ***Kuboniwa et al. (1995***) (Fig. 5). During the Ca^2+^ binding and conformational transition, the hydrophobic cluster formed by these residues, including Phe16, Phe19, Ile27, Leu32, Val35, Ile52, Val55, Ile63 and Phe68, are almost rigid (Fig. S5). Comparison of the structures in the apo- and holo-states suggests that the two helices of the EF-hand rotate around the hydrophobic cluster during the conformational transition.

**Figure 5.**
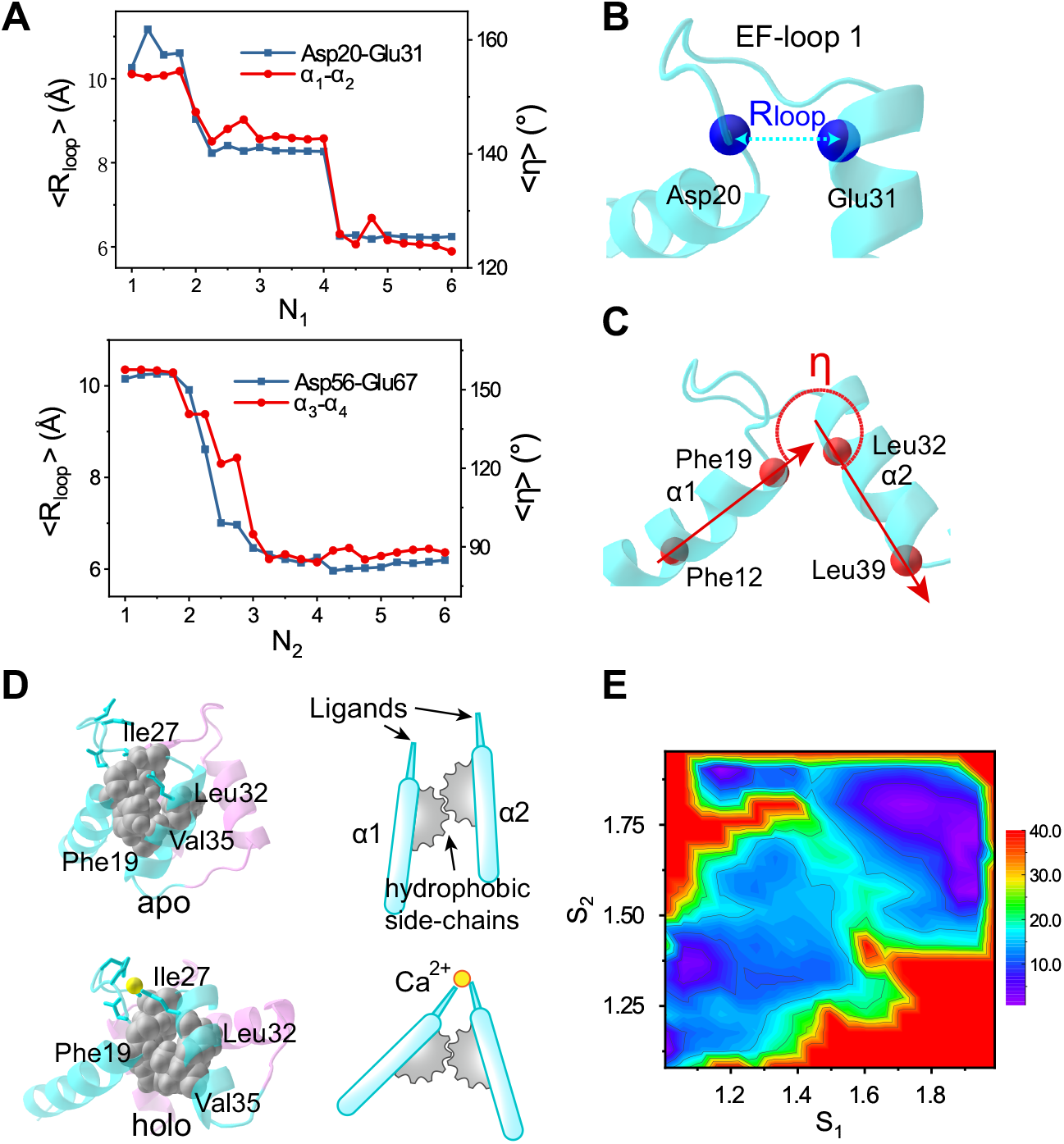
Ca^2+^ binding induced rotation of EF-hand helices. (A) The distance between close ends of EF-loop 1 (represented by the C_*α*_ atoms from Asp20 and Glu31) and inter-helical angle between the helices in EF_1_ as a function of ***N***_1_ (upper panel). The same results for the EF_2_ are shown in the lower panel of (A). (B) Schematic diagram illustrating the end-end distance of EF-loop1 and (C) the inter-helical angles of EF_1_. The end-end distance of EF-loop is represented by the distance between C_*α*_ atoms: EF-loop1 by Asp20–Glu31, and EF-loop2 by Asp56–Glu67. The direction of helices are represented by vectors pointing from one residue’s C_*α*_ atom to another residue’s C_*α*_ atom: *α*1 by Phe12–Phe19, *α*2 by Glu31–Leu39, *α*3 by Leu48–Val55, and *α*4 by Phe68–Lys75. (D) Schematic diagram of the Ca^2+^ binding induced EF-hand helix rotation. Black spheres indicate the hydrophobic core in EF_1_. (E) Free energy profile of nCaM conformational transition projected onto ***S***_1_ and ***S***_2_. The free energy scale is in kJ/mol.

To demonstrate the coupling between the Ca^2+^ binding and the EF-hand opening, we calculated the distance between the EF-loop ends (*R*_*loop*_, is the distance between the blue residues in Fig. 5B) and the inter-helical angles between the helices (*η*, denoted by the red arrows in Fig. 5C, which are represented by vectors pointing from one residue’s C_*α*_ to another) of the two EF-hands, respectively. The results in Fig. 5A show that *R*_*loop*_ decreases upon Ca^2+^ binding, thus pulling the ends of the EF-hand helices close together (blue lines). In particular, coordination of the fourth (third) ligand to Ca^2+^, which corresponds to the binding of the terminal residues of the EF-hand loops to the Ca^2+^ by water bridged interactions (direct interactions) in EF_1_ (EF_2_), has the most prominent effect in changing the distance between the two close ends. Meanwhile, the angles between the two helices increase (red lines), indicating the rotation of the EF-hand helices. In the process of rotation, the hydrophobic cluster, which is stable during the Ca^2+^ binding, acts as the pivot (Fig. 5D).

The results described above show that the large conformational change in the two EF-hand motif occurs at different stages of the coordination (Fig. 5A and Fig. 2). Analysis of the correlation between the coordination processes of the two EF-hand motifs demonstrated that the Ca^2+^ binding proceeds earlier in the EF_1_ than that in the EF_2_ (Fig. S6). When ***N***_1_ = 3 or 4, the ***N***_2_ dominantly takes the values of 1 or 2, and the EF_2_ mainly stays at the closed conformation. As shown in Fig. 5E, the conformational changes in the two EF-hands mostly follow the diagonal line of the two dimensional free energy landscape, which suggests tight coupling and cooperativity between the two EF-hands. When ***N***_2_ = 3 or 4, the native coordination bonds of the EF_1_ are almost fully formed, which favors a open-like conformation for the EF_2_ due to the tight coupling between the two EF-hand motifs, even though the resulting native coordination number of the EF_2_ is relatively small. The tight coupling between the two EF-hands is consistent with previous experimental observations ***Linse et al. (1991***); ***Malmendal et al. (1999***).

Taken together, these results demonstrate that the rotation of the EF-hand helices is coupled to Ca^2+^ binding, which is consistent with the free energy landscapes shown in Fig. 2. Ca^2+^ binding induced rotation of the EF-hand helices was also noted in a simulation study ***Goto et al. (2004***).

### Bridging water molecules lower the free energy barrier between the allosteric states

As mentioned above, water-bridged interactions play a crucial role during Ca^2+^ binding, and hence in the allosteric transitions in CaM. To further demonstrate the role of water molecules, we performed detailed analyses of the ensemble of conformations containing water-bridged coordinated structures. We assume that a water-bridged coordination is formed if the following conditions are satisfied: (i) the distance from oxygen of water molecule (*O*_***w***_) to Ca^2+^ is less than 2.8Å; and (ii) the hydrogen bond is formed between water and the native ligand. For each residue listed in Table 1, the Ca^2+^-water-native ligand conformation exists at certain stages of the binding process with high probability. These results suggest that before the final chelation process, the native ligands already develop indirect interactions with Ca^2+^ mediated by water molecules. For example, after binding of Ca^2+^ to five of the native ligands of EF_2_, the last native ligand, Asn60 or Thr62, is not free”, but has a probability of 0.65 to be bridged to the Ca^2+^ by a water molecule (Table 1).

To quantitatively evaluate the contribution of water molecules to the Ca^2+^ binding process, we characterize the binding involving water bridge in Glu31 to the Ca^2+^ in the final step of the Ca^2+^ coordination to EF_1_. We used well-tempered metadynamics to extract the free energy profile along the collective variable ***D***, which is defined as the distance between Ca^2+^ and the closest side-chain oxygen of Glu31 (**Methods and Materials**). As a control, we also designed a system in which the electrostatic interactions between the bridging water molecule and Glu31 side-chain atoms were turned off. The differences between the two simulations reflect the net effect of the bridging water on the binding free energy landscape. When the interaction of the bridging water includes the electrostatic potential, an intermediate is present in the free energy profile at ***D*** *≈* 4.3Å (see Fig. 6A, black line), in which the Glu31 is bridged to the Ca^2+^ by the bridging water molecule (Fig. 6B), leading to a low free energy barrier for ion coordination. In contrast, when the electrostatic interactions involving the bridging water molecule are switched off, a high free energy barrier is observed (∼ 6kJ/mol, shown in Fig. 6A, red line). We surmise that water reduces the free energy barrier by bridging the Ca^2+^–Glu31 interaction, therefore speeding up the coordination between the Ca^2+^ and Glu31.

**Figure 6.**
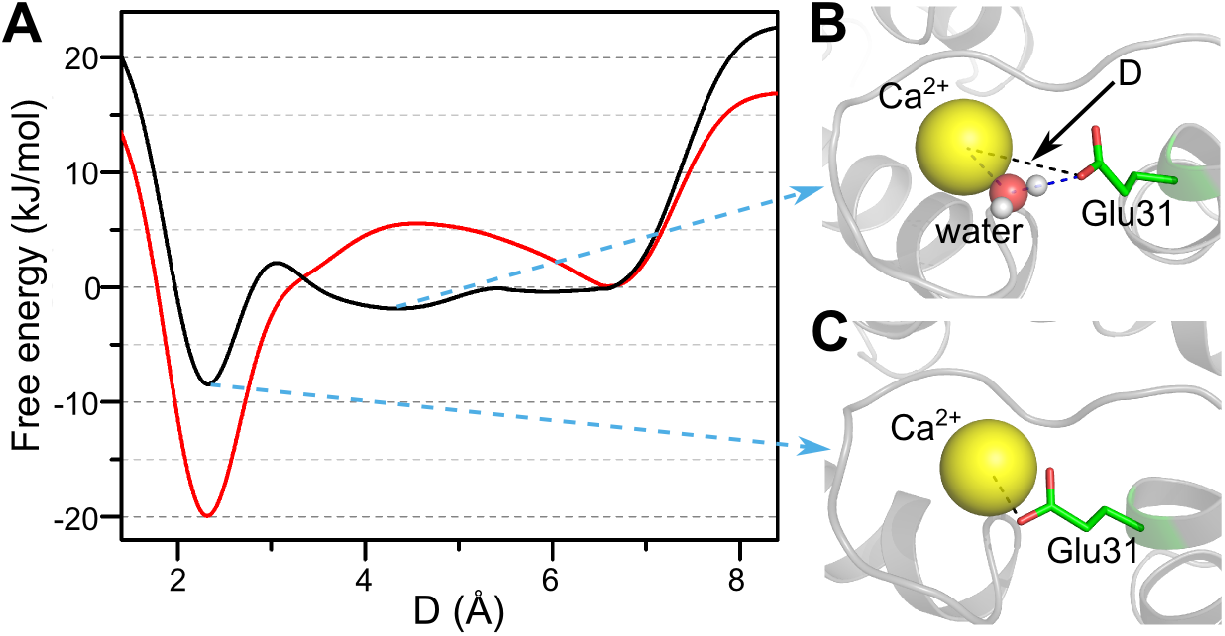
Free energy profiles of water bridged coordination of Glu31 side-chain oxygen atoms to the Ca^2+^. (A) Potential of mean force for the binding/unbinding process of Glu31 to Ca^2+^ as a function of the minimum distance between side-chain oxygen atoms of Glu31 and Ca^2+^. The black line illustrates the results with the water molecules being realistically treated. The red line shows the results with electrostatic interactions involving the bridging water molecules being turned off. (B) The water-bridged coordination of Glu31 oxygen and Ca^2+^, where ***D*** = 4.3Å is a local free energy minimum. (C) shows the direct coordination bond of Glu31 oxygen and the Ca^2+^, which corresponds to the most stable state at ***D*** = 2.3Å.

Two possible effects may contribute to the reduction of the free energy barrier by the bridging water. First, the bridging waters effectively introduce attractive interactions between Ca^2+^ and ligands, which tends to decrease the free energy barrier and therefore reduce ruggedness of ligand exchange landscape by acting as a lubricant. Second, the bridging water reduces the translational entropy of the captured Ca^2+^, therefore decreasing the entropy penalty during the Ca^2+^–Glu31 coordination, which could also contribute to the reduction of the free energy barrier. We propose that the contribution of water molecule to the binding of metal ions should be a common feature in other metal-ion induced biological processes.

## Conclusions

Allosteric transitions are ubiquitous in biology ***Thirumalai et al. (2019***). Nowhere is it more prominent in modulating large conformational changes over long distances (nearly 20Å) than it is in CaM induced by Ca^2+^ binding. Using all-atom molecular dynamics simulations in conjunction with metadynamics, which enables efficient sampling of the conformational space, we have established that in the Ca^2+^ binding coupled allosteric transition process water-bridged interactions play crucial role. In addition to participating in the native coordination of Ca^2+^ as widely observed in the crystal structures of metalloproteins ***Andreini et al. (2012***), the transient bridging water molecules between Ca^2+^ and binding residues during the Ca^2+^ binding/unbinding process tend to reduce the ruggedness of ligand exchange landscape by acting as a lubricant, facilitating the Ca^2+^ coupled protein allosteric motions. The high free energy cost dehydration of Ca^2+^ occurs in steps with the number of waters lost at different stages being compensated by gain in chelation to the charged residues in the loops of the EF-hands. The overall picture that emerges is that there is a coordinated interplay between ligand binding, loss of water around the cation, and subsequent changes in the conformations of the protein. We propose that similar mechanisms, with water playing an important role, might also mediate allosteric transitions in molecular machines, such as molecular chaperones, molecular motors, and cell adhesion molecules, in which allosteric transitions are often triggered by cations. The ability to undergo dehydration depends on ion charge density, which for Ca^2+^ is in the optimal range ***Lee et al. (2017***). Our results also demonstrated that there is an asymmetry in the Ca^2+^ binding and conformational transitions in the EF-hand pair of isolated CaM domains despite the overall structural similarity. The Ca^2+^-coupled closed to open transitions in the two EF-hand motifs occur by an entirely different mechanism, which could facilitate the differing functional requirements of CaM.

A key event in the Ca^2+^ coupled allostery of CaM is the dehydration of Ca^2+^. Had the charge density been even somewhat larger, as is the case in Mg^2+^ the dehydration would not occur rapidly enough to induce the functionally required allosteric transition. Thus, the ease of fast dehydration in Ca^2+^ may well have been a consequence of evolution in CaM and other Ca^2+^ systems, such as the lever arm in myosin motors.

## Methods and Materials

### Molecular dynamics simulations

We used the NMR structure of *Xenopus* nCaM for the holo form with 1J7O ***Kuboniwa et al. (1995***) as the Protein Data Bank (PDB) entry. The apo form with the same sequence (residue 1 to 76) was taken from the NMR structure of *X. laevis* CaM with PDB code 1CFD ***Chou et al. (2001***). All MD simulations were carried out using GROMACS 2019.6 ***Abraham et al. (2015***) with the OPLS/AA force field ***Kaminski et al. (2001***). The Ca^2+^-nCaM system was solvated in the SPC/E water box together with Na^+^ and Cl^−^ ions to mimic the ion concentration of 0.15***M***. The whole system had 15,117 atoms, of which 13,929 atoms were from water. We used the periodic boundary condition where the size of unit box is ∼ 49.2 × 55.9 × 56.7 Å^3^. Energy minimization was performed before the MD simulations. Before running metadynamics simulations the Ca^2+^-nCaM systems were heated to 300K in the NVT ensemble and then equilibrated for 10 ns MD under NPT conditions, with ***P*** = 1atm and ***T*** = 300K. Periodic conditions were used and electrostatic interactions were calculated using the particle-mesh Ewald algorithm. The simulations for the cCaM were conduced similarly.

### Collective Variables (CVs) in metadynamics

We used the “bias-exchange” ***Piana and Laio (2007***) form and “well-tempered” ***Barducci et al. (2008***) algorithm of metadynamics to accelerate conformational sampling, using the PLUMED 2.6.1 package ***Tribello et al. (2014***). The CVs used to study the Ca^2+^-binding coupled conformational change of CaM were chosen to describe the rate-limiting events involving the binding of Ca^2+^ and the subsequent conformational transition of the protein.

The events driving the closed to open transition involve movement of a number of particles from distant parts of CaM domains. In order to capture such an allosteric movement, we need suitable reaction coordinates. Since the major events in the closed → open transition are triggered by Ca^2+^ binding, we constructed CV involving the oxygen atoms (ligands) of the residues in the holo structure that are coordinated to Ca^2+^. In terms of these native ligands, we define the collective variables, ***N***_*α*_ (*α* = 1 and 2 (3 and 4) for EF_1_ and EF_2_ (EF_3_ and EF_4_), respectively, of the nCaM and cCaM) to describe the number of native ligands that are coordinated to Ca^2+^:

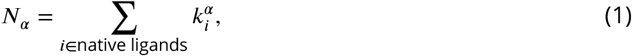

where 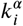 is defined using the switching function,

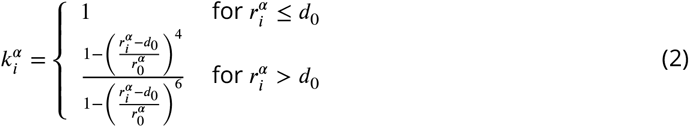

In equation (2), 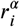 is the distance from the *i*^*th*^ native ligand to the *α*^*th*^ Ca^2+^. By native ligand we mean the oxygen atoms from distinct amino acids that are linked to Ca^2+^ and are present in the native (NMR) holo structure. We chose *d*_0_ = 2.5Å and *r*_0_ = 0.02Å. For all the Asp residues, the two oxygens from the same residue were considered “degenerate”, namely, when more than one oxygen is coordinated to Ca^2+^, we consider only one of them to be a native ligand. However, for the Glu residues in the 12^*th*^ position of the EF-loops of nCaM and cCaM, two oxygens are treated as individual ligands. We treated coordination of Ca^2+^ to Asp differently from binding to Glu because in the holo structure of the nCaM and cCaM (PDB entry 1J7O and 1J7P, respectively) Ca^2+^ is coordinated to both oxygen atoms of Glu but only to one of the Asp oxygen atoms.

We also used the “path collective variables” calculated using,

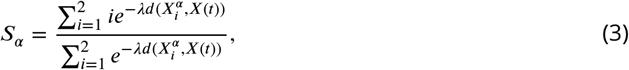

where *α* = 1, 2, 3, and 4 dictates the *α*^*th*^ EF-hand, and 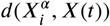 is the mean square deviation from the closed (*i* = 1) and open (*i* = 2) structure for a given conformation *X*(*t*). A small value of ***S***_*α*_ corresponds to closed structure and a large value of ***S***_*α*_ corresponds to open structure. We used *λ* = 20 in our simulations.

To use the bias-exchange method with metadynamics, we designed 5 replicas. In the first replica, there is no bias in the CVs. In each of the other four replicas, a bias was applied to one of CVs, including ***N***_1_, ***N***_2_, ***S***_1_, and ***S***_2_ for the nCaM simulations, and ***N***_3_,***N***_4_, ***S***_3_, and ***S***_4_ for the cCaM simulations. Exchange between replicas was attempted every 10,000 MD steps, and each replica lasts for 800ns. In the well-tempered algorithm, we used a bias factor of 200 for the rescaling of the Gaussian height, and 0.3kJ/mol as the initial Gaussian height. More detailed descriptions of the simulation procedure can be found in Supplementary Methods and Supplementary Fig.S7, S8 and S9 online.

### Water-bridged coordination of Glu31 to the Ca^2+^

We also used well-tempered metadynamics to study water-bridged coordination of Glu31 to the Ca^2+^ for the nCaM. The minimum distance between the Ca^2+^ and two oxygen atoms of Glu31 is chosen as the collective variable ***D Tribello et al. (2014***), which we define as:

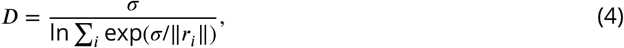

where *r*_*i*_ (*i* = 1, 2) is the distance between each oxygen and the Ca^2+^, and *σ* = 5000Å. The well-tempered rescaling factor is 12, and the initial Gaussian height is 0.2kJ/mol.

In the control simulation, once a water molecule comes into the first ligand shell of Ca^2+^ and simultaneously within 3.5Å distance of any of the two oxygen atoms of Glu31, the electrostatic interactions between the water and the side-chain heavy atoms of Glu31 are turned off. This was accomplished by modifying the source code of GROMACS package.

To achieve better convergence we use a restraining potential at ***D*** = 6.5Å to limit the region of the phase space accessible during the simulation:

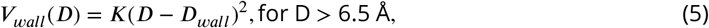

where ***K*** = 50*k****J*** /*mol*/Å^2^ and ***D***_*wall*_ = 6.5Å. We performed 20 independent 100ns simulations for each system, and then calculated the ensemble average to obtain the free energy reported in Fig. 6. Comparison of the simulations with and without electrostatic interactions allows us to assess the role water plays in modulating the binding of Ca^2+^.

## Acknowledgments

This work was supported by the National Natural Science Foundation of China (Nos. 11974173, 11934008). Part of this work was done while DT was a visiting Professor in Nanjing University. DT acknowledges the NSF (CHE09–14033) for partial support of this work. The computing resources were provided by the High Performance Computing Center of Nanjing University.

## Supplementary information

### Supplementary Methods

#### Well-tempered Metadynamics algorithm

The metadynamics algorithm was used to enhance conformational sampling in simulations so that Free Energy Surfaces (FES) could be reliably computed. In complex systems, it is necessary to identify physically reasonable collective variables (CVs) to describe large scale conformational transitions. In the canonical metadynamics algorithm, the system is biased through the addition of a sum of repulsive Gaussian terms to the system potential energy function. The biasing potentials are usually applied to the CVs. The total biasing potential applied on a set of CV, *Y* (*x*) = (*Y*_1_(*x*), *Y*_2_(*x*), …, *Y*_*n*_(*x*)), at time *t* during the simulation can be expressed as [1]:

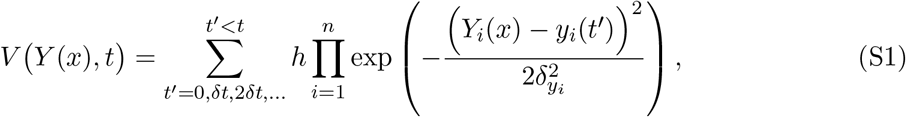

where *h* is the height of each Gaussian potential, 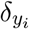 is the Gaussian width associated with the CV *Y*_*i*_(*x*), *δt* determines the time step at which the biasing potentials are added. In order to accelerate the convergence of metadynamics, the well-tempered algorithm was designed [2], in which the height of the Gaussian potentials is systematically attenuated during the course of the simulations using 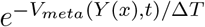, where Δ*T* represents a characteristic energy [2]. The form of the biasing potential used in our simulations is

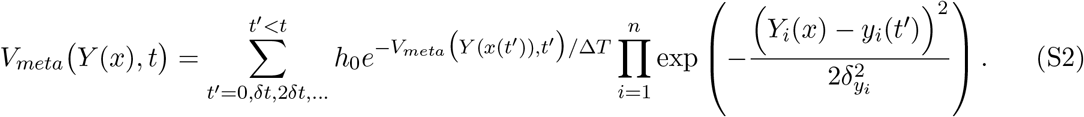

where *h*_0_ is the initial Gaussian height. In practice, the rate of decrease is controlled by the bias factor *η* = (*T* + Δ*T*)*/T* [3], where *T* is the system temperature. Combining the replica exchange methodology and metadynamics, the bias-exchange metadynamics was designed to enhance the sampling efficiency[4]. As an inherent feature of the algorithm, the converged metadynamics runs enable the calculation of the canonical probability distribution of the CVs directly. However, the statistics of other degrees of freedom is affected by the bias, and require “reweighting” techniques to reconstruct the unbiased canonical probability distribution[5]. In particular, for well-tempered metadynamics, the reconstruction algorithm is described in Ref.[6], and can be found in the PLUMED package [7], which is used in our work.

#### Technical details of the simulations

During the metadynamics simulations, we applied restraining potentials to the values of the MSD (mean square deviation) of a conformation with respect to the folded states to limit the accessible region of phase space sampled by the protein. The use of such restraining potentials physically prevents unreasonable unfolding of nCaM. The functional form of the restraining potential is [3]:

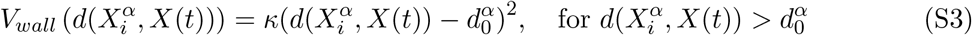

where the energy constant *κ* = 0.8kJ/Å^4^, 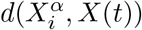 is the MSD from the closed (*i* = 1) or open (*i* = 2) structure of the *α*^*th*^ (*α* = 1, 2) EF-hand for a given conformation *X*(*t*), the limit 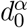 (*α* = 1, 2) is chosen as 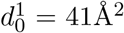 for EF_1_ and 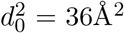 for EF_2_. The terminal residues with high flexibility were not included in the calculations of the MSD values. For better convergence, we prevent Ca^2+^ ions from leaving too far from the binding sites by applying a lower limit to the coordination number, as one of the reaction coordinates.

#### Convergence of metadynamics simulations

We sampled the solution conformations of nCaM and the associated Ca^2+^ binding using well-tempered bias-exchange metadynamics with 5 replicas (in four of which biasing potentials were applied). The simulations were initiated from random conformations, and were continued for 800ns. We assessed the convergence of the metadynamics simulations by ensuring that the free-energy differences between all the major populated regions for each CV reach a steady state. For the four biased replicas, this condition was satisfied after about 400ns of sampling (see Fig.S7). In addition, as an embedded property of well-tempered metadynamics, the automatically rescaled Gaussian height guarantees convergence, and avoids fluctuations around the correct free energy value [2, 8]. Therefore, we checked the height of the biasing Gaussian potentials during the simulations. As can be seen in Fig.S8, 200ns after sampling starts, all the Gaussians are less than 1/10 of the initial height, which also indicates the convergence of our metadynamics simulations. In addition, we analyzed the evolution of two dimensional FESs during the simulations. As an example, FESs of *N*_1_-*S*_1_ at different simulation times are shown in Fig.S9. The evolution of FESs also show that they are converged after 400ns of simulations. Thus, we conclude that the calculated FESs as a function of CVs are well converged in our simulations.

#### Coordination probability

Binding pathways are quantified using the probability, 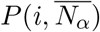, that the *i*^*th*^ residue is coordinated to Ca^2+^ when the total ligand number of EF-hand *α* is 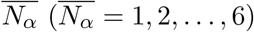. We define 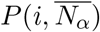 using

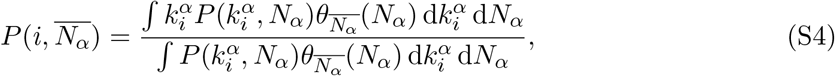

where *N*_*α*_ and 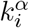 are defined in equation (1) and equation (2) (in the main text), respectively, and 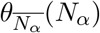 is a step function:

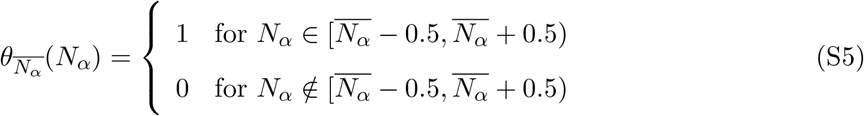

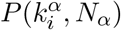 is a two dimensional probability distribution of the variables 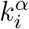 and *N*_*α*_, which cannot be directly calculated as the average over time from the metadynamics. We use the reweighting techniques described in Ref.[6] to reconstruct the unbiased distribution. We calculated the two-dimensional probability distributions on a 60 *×* 60 grid. Then we constructed bins on the *N*_*α*_ dimension as (0, 0.5), [0.5, 1.5), [1.5, 2.5), …, [4.5, 5.5), [5.5, 6], and binned any 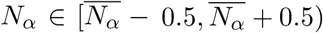 into integer values 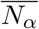.

The occurrence probability of the water-bridged coordination has the same form as equation (S4), except for the definition of *k*_*i*_. For the water-bridged coordination, *k*_*i*_ = 1 when the oxygen of water is within 2.8Å of the Ca^2+^ and at the same time within 3.5Å of the oxygens from a native residue involved in ligand formation, and is zero otherwise.

## Supplementary Figures

**Figure S1:**
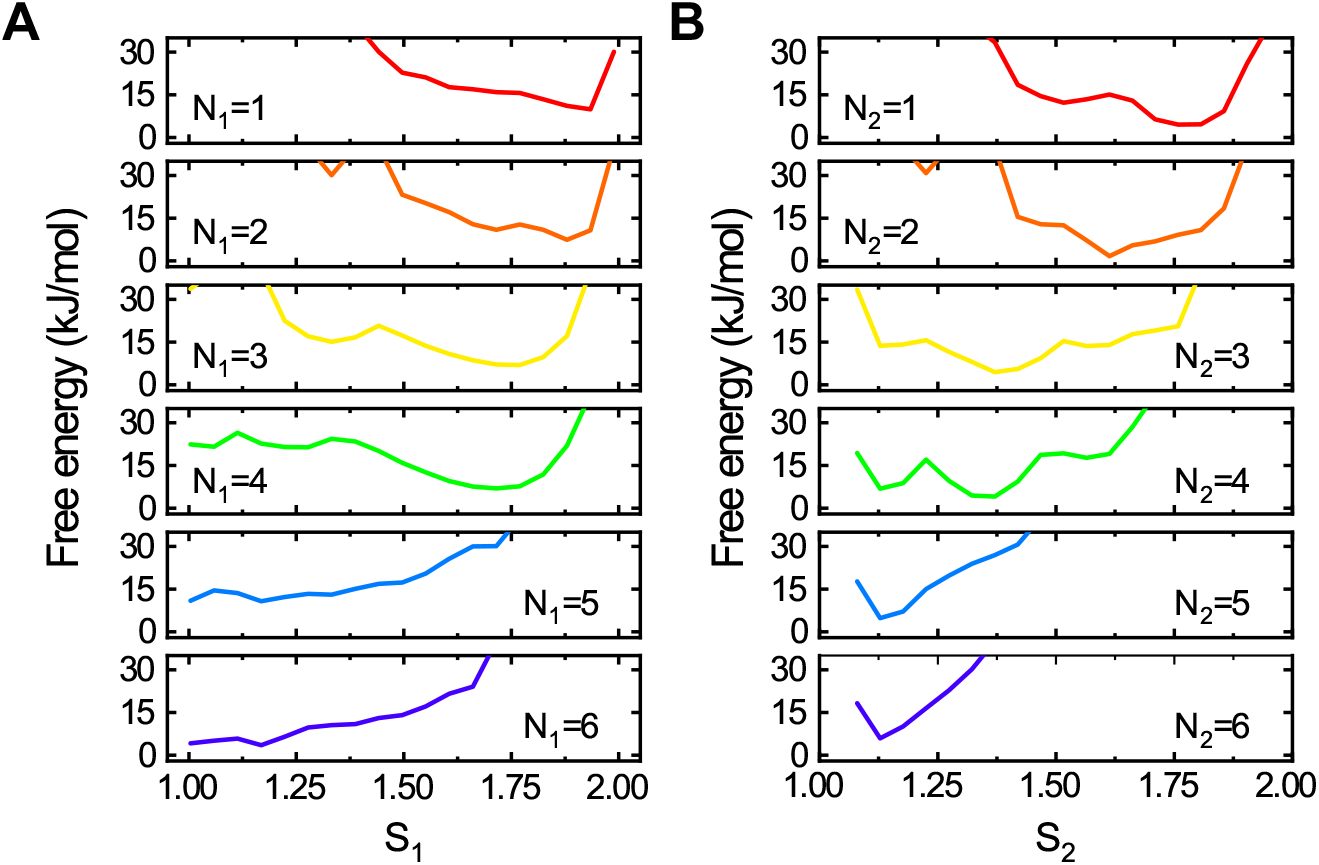
One-dimensional free energy profiles of the Ca^2+^ coupled conformational transitions of EF_1_ (A) and EF_2_ (B) along the collective variables *S*_*α*_ (*α*=1 and 2 for the EF_1_ and EF_2_, respectively) at different binding stages described by *N*_*α*_, which is the number of native ligands that are coordinated to Ca^2+^.

**Figure S2:**
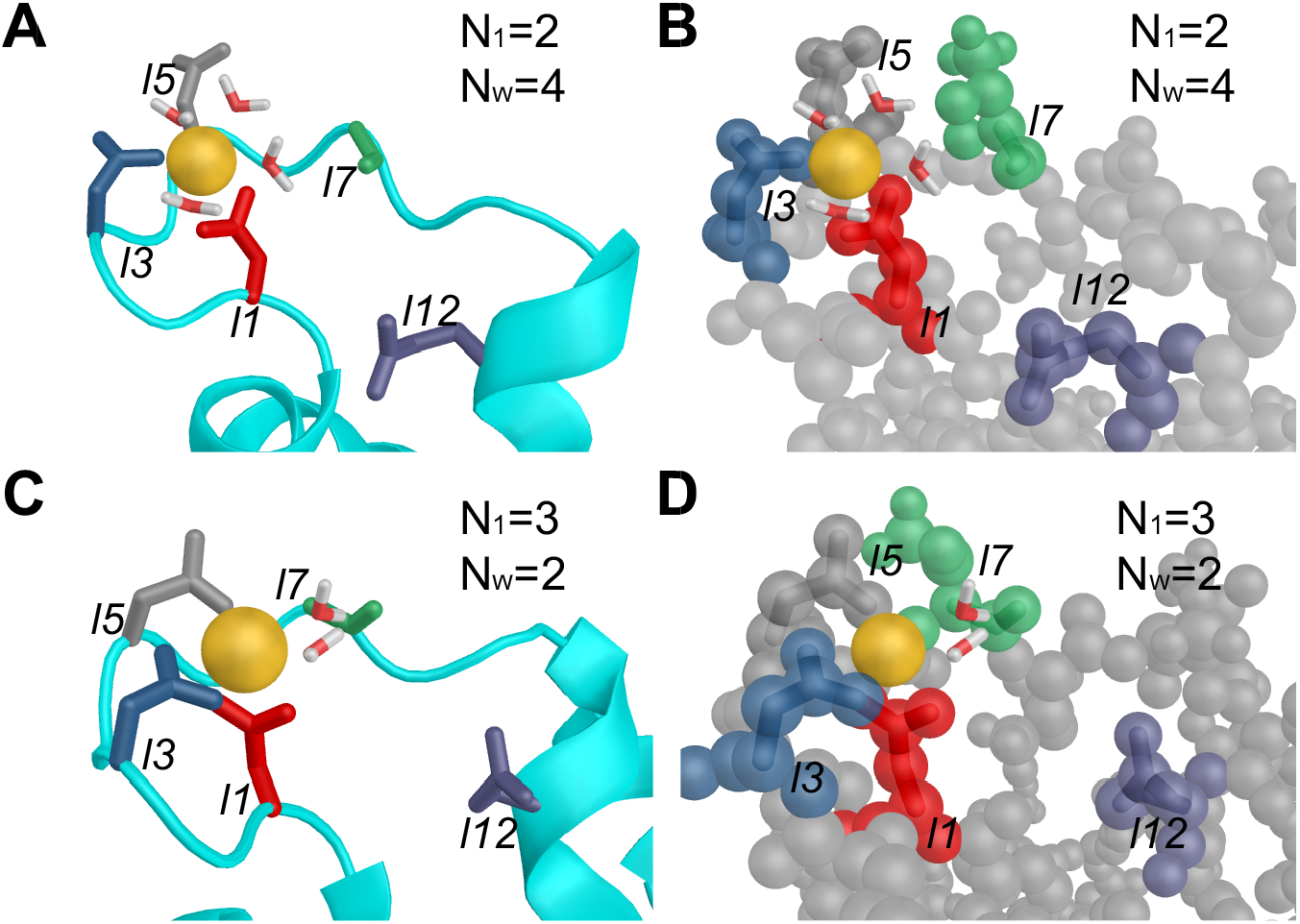
Dehydration of Ca^2+^ is coupled to the binding of native ligands from *N*_1_ = 2 to *N*_1_ = 3 in EF_1_. (A) and (B) show the coordination conformation of Ca^2+^ when *N*_1_ = 2, using “new cartoon” (A) and “sphere” (B) drawing methods. *N*_*w*_ = 4 represents that four water molecules are coordinated to Ca^2+^. (C) and (D) show the coordination of Ca^2+^ when *N*_1_ = 3 and *N*_*w*_ = 2. These figures illustrate the decrease of *N*_*w*_ by two when the third native ligand is bound to Ca^2+^. Possible reasons are the potential coordination of two side-chain oxygens from one aspartate to Ca^2+^, such as Asp20 (*l*1) in (C), and volume exclusion effects at the binding site in protein, as shown in (D).

**Figure S3:**
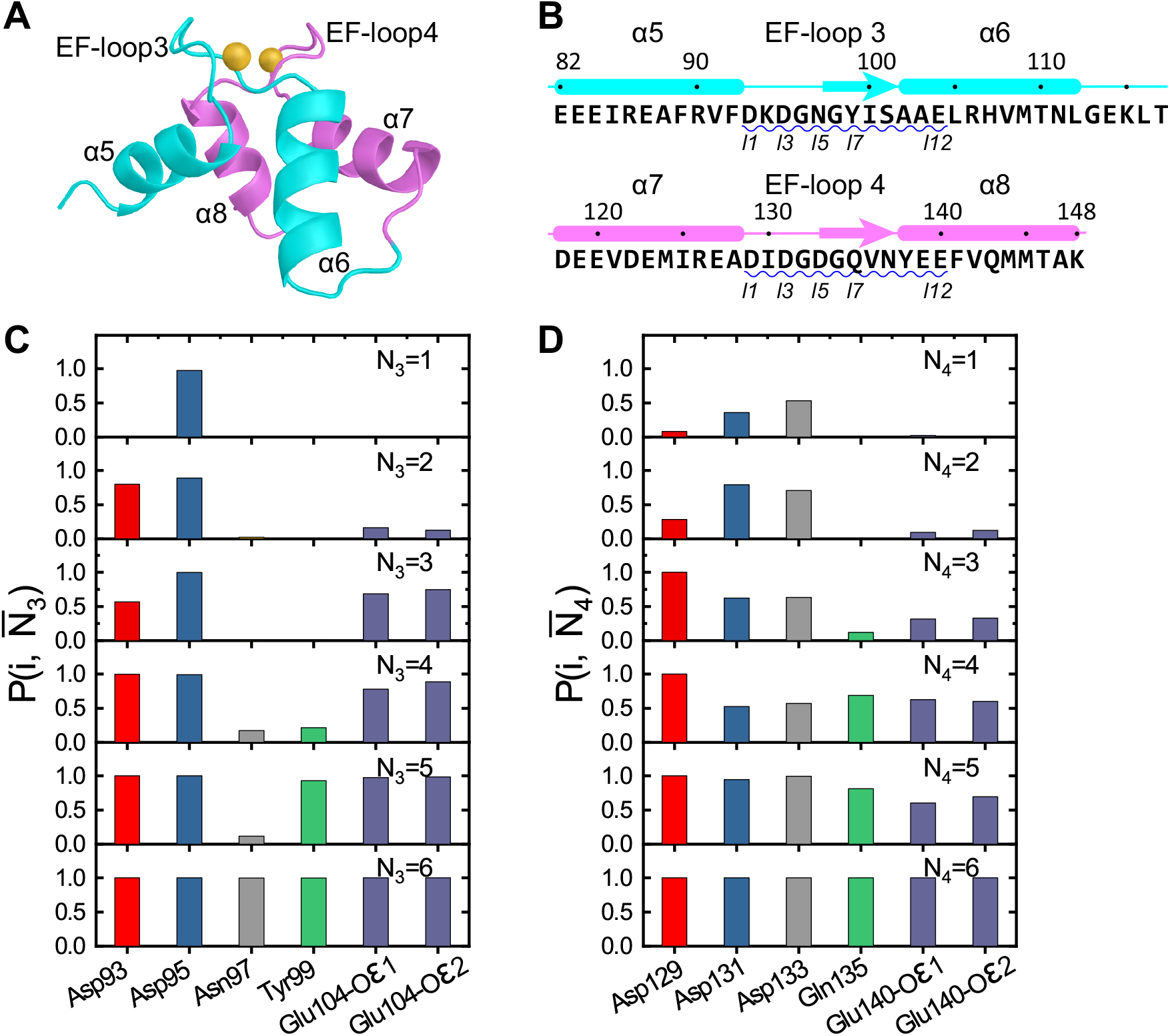
Ca^2+^ binding coupled to conformational changes of the EF-hand motifs in cCaM and molecular mechanism of the Ca^2+^ binding. (A)Cartoon representation of the three-dimensional structure of the cCaM in the holo state. (B) Sequence of cCaM and corresponding secondary structures. (C) Coordination probability 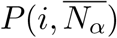 for the six native liganding residues of EF_3_. Different liganding residues are colored in different colors. (D) Same as C but for EF_4_.

**Figure S4:**
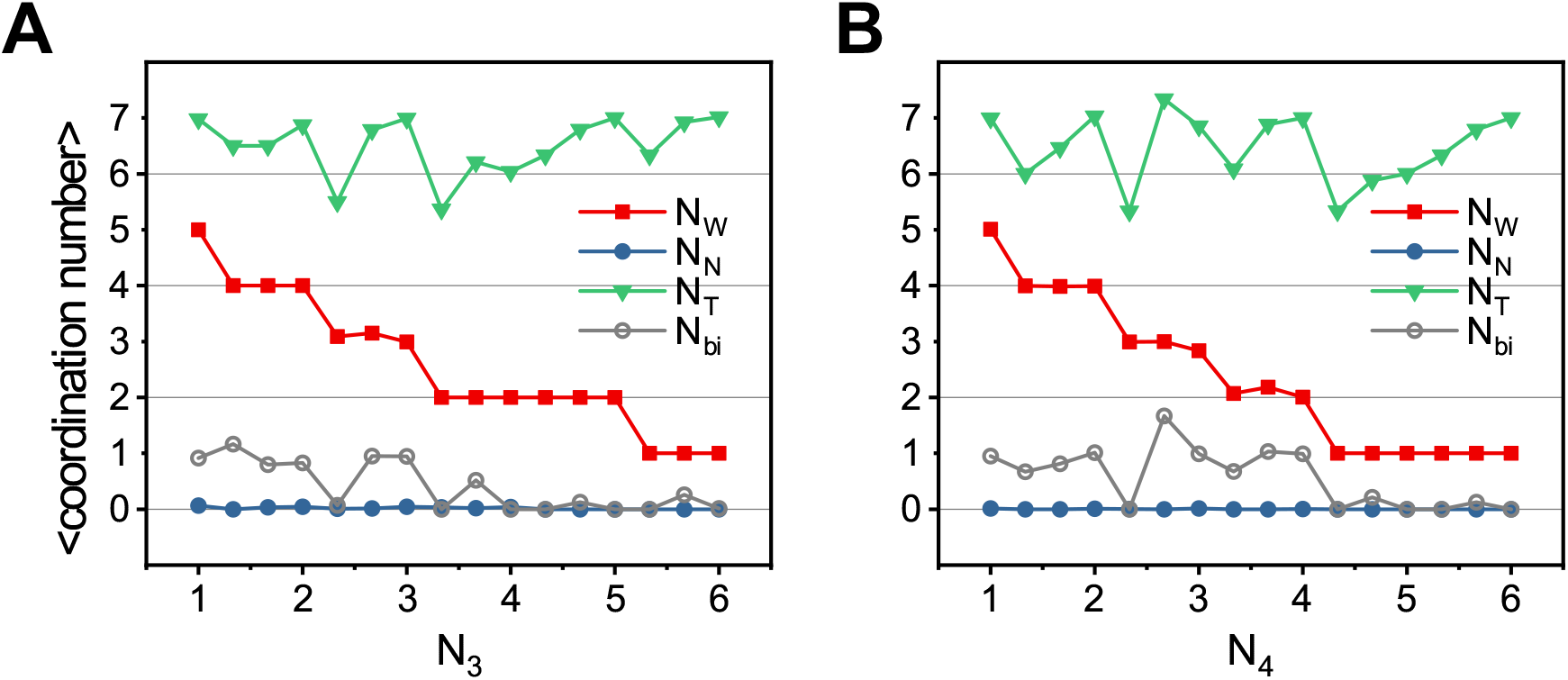
Number of coordinated water molecules (*N*_*W*_), non-native ligands (*N*_*N*_) as well as the total coordination number (*N*_*T*_) as functions of *N*_3_ and *N*_4_ for EF_3_ (A) and EF_4_ (B). Although only one carboxyl oxygen atom is coordinated to the Ca^2+^ at the native coordination structure, the second carboxyl oxygen atom may also bind to the Ca^2+^ in the intermediate state of the coordination. The average number of such non-native bidentate ligand was also shown in panels A and B (N_*bi*_, gray).

**Figure S5:**
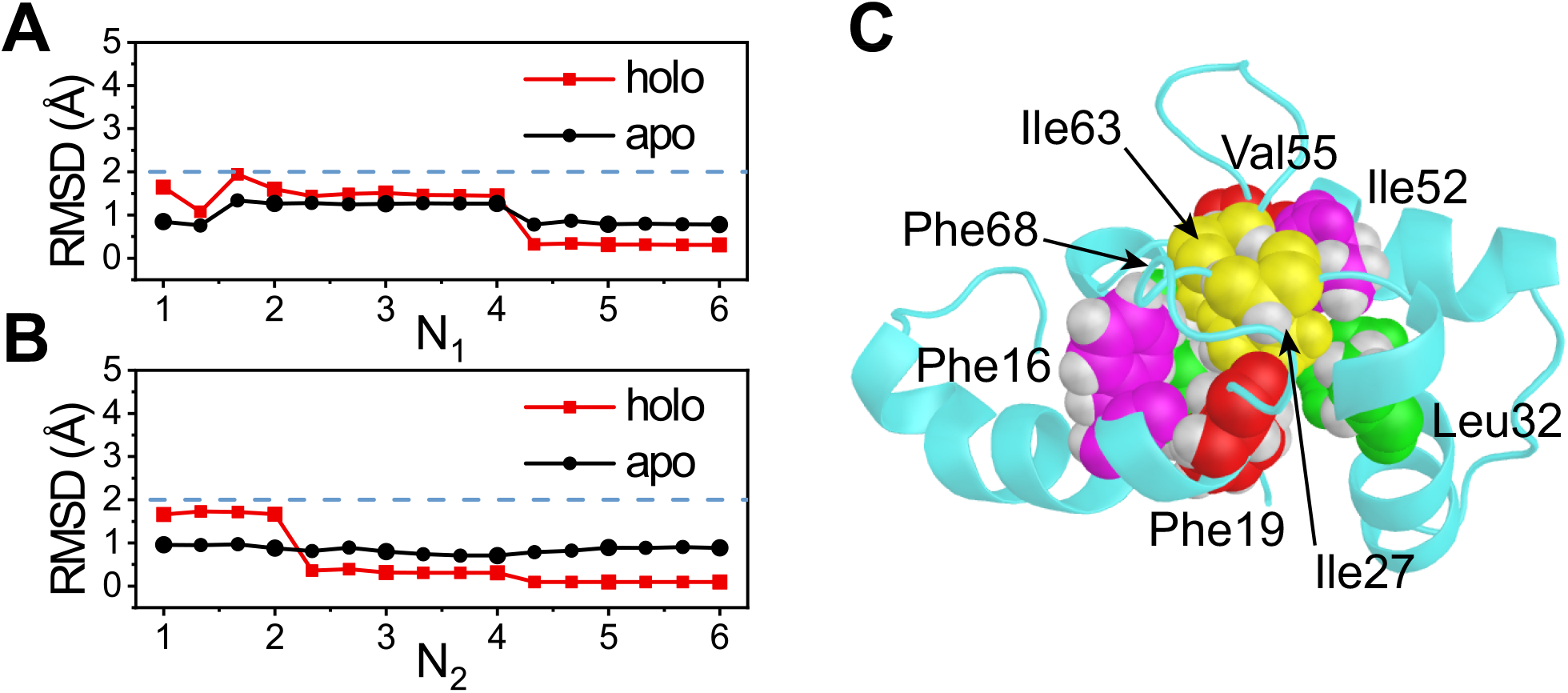
Allosteric conformational changes in the hydrophobic cluster of nCaM. (A, B) The RMSD of the hydrophobic residues at conserved positions for the two EF-hand motifs as a function of *N*_1_ (A) and *N*_2_ (B). The relevant residues in the hydrophobic core are Phe16, Phe19, Ile27, Leu32 for EF_1_, and Ile52, Val55, Ile63, Phe68 for EF_2_. (C) The hydrophobic cluster in the nCaM shows that it is densely packed.

**Figure S6:**
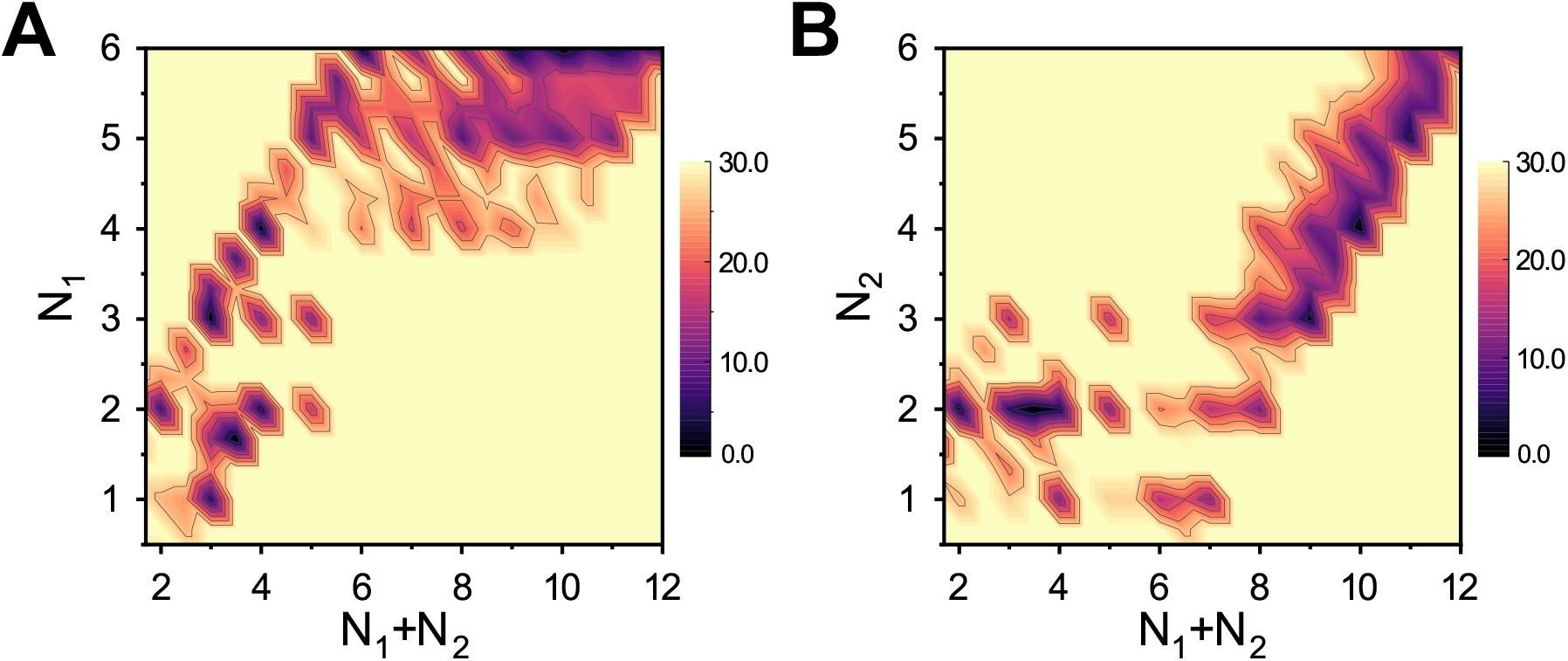
Two-dimensional free energy profiles illustrating the coupling of Ca^2+^ coordinations between EF1 and EF2. (A, B) Free energy profiles projected onto the collective variables (*N*_1_+ *N*_2_, *N*_1_) (A) and (*N*_1_ + *N*_2_, *S*_2_) (B), respectively. The free energy scales are in kJ/mol.

**Figure S7:**
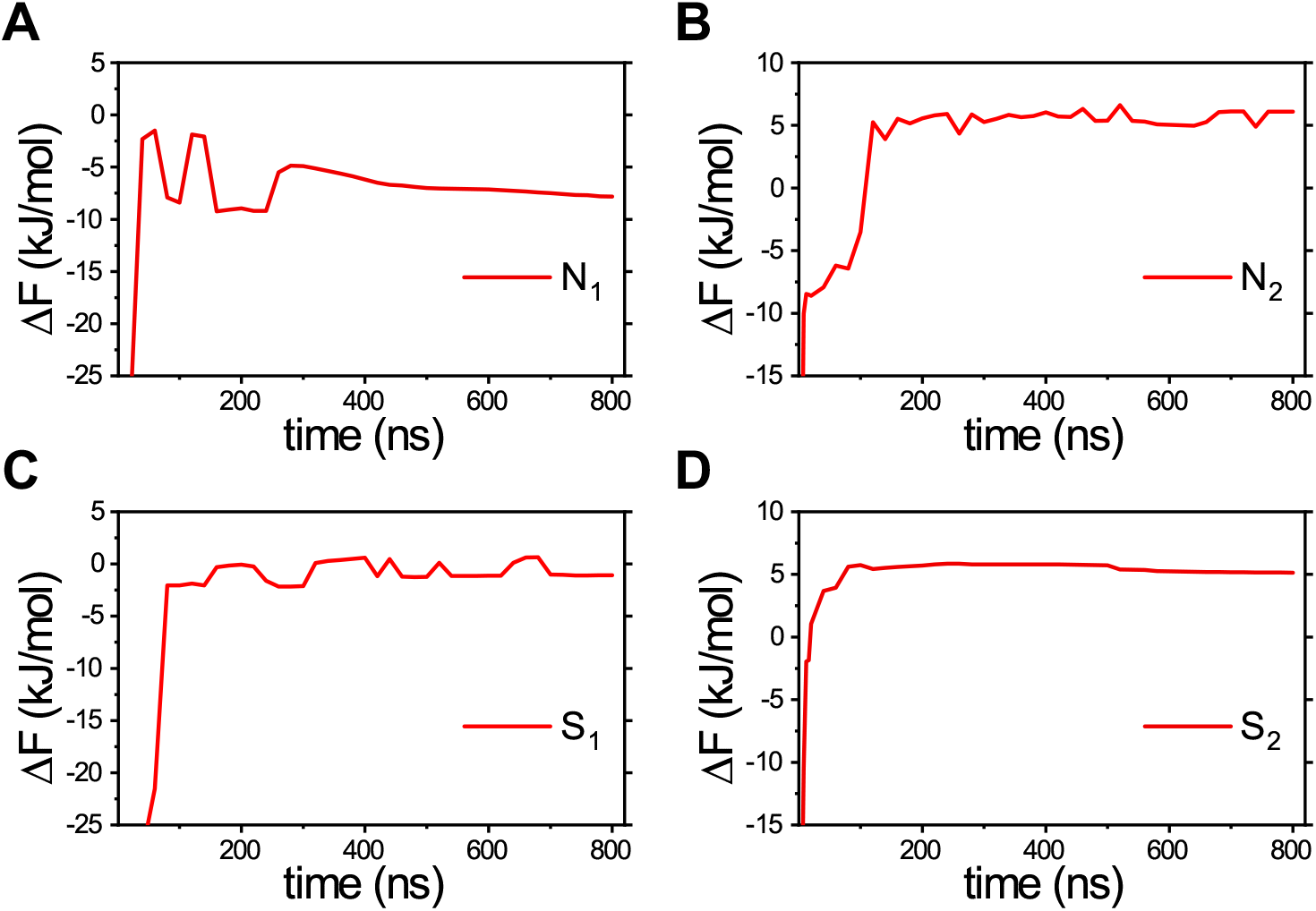
The free energy difference between local minima in a one dimensional FES corresponding to a specific CV as a function of the simulation time. (A) and (B) Free energy differences between the regions at *N*_*α*_ = 6 and *N*_*α*_ = 2 for EF_1_ and EF_2_ (Δ*F* = *F* (*N*_*α*_ = 6) −*F* (*N*_*α*_ = 2), *α* = 1, 2). (C) and (D) Free energy differences between the open conformation and the closed conformation for the two EF-hands (Δ*F* = *F* (*S*_*α*_ *<* 1.2) − *F* (*S*_*α*_ *>* 1.8), *α* = 1, 2). In (A-D) the Δ*F* s reach steady values after ∼300 ns of simulations, which indicate the convergence of metadynamics simulations.

**Figure S8:**
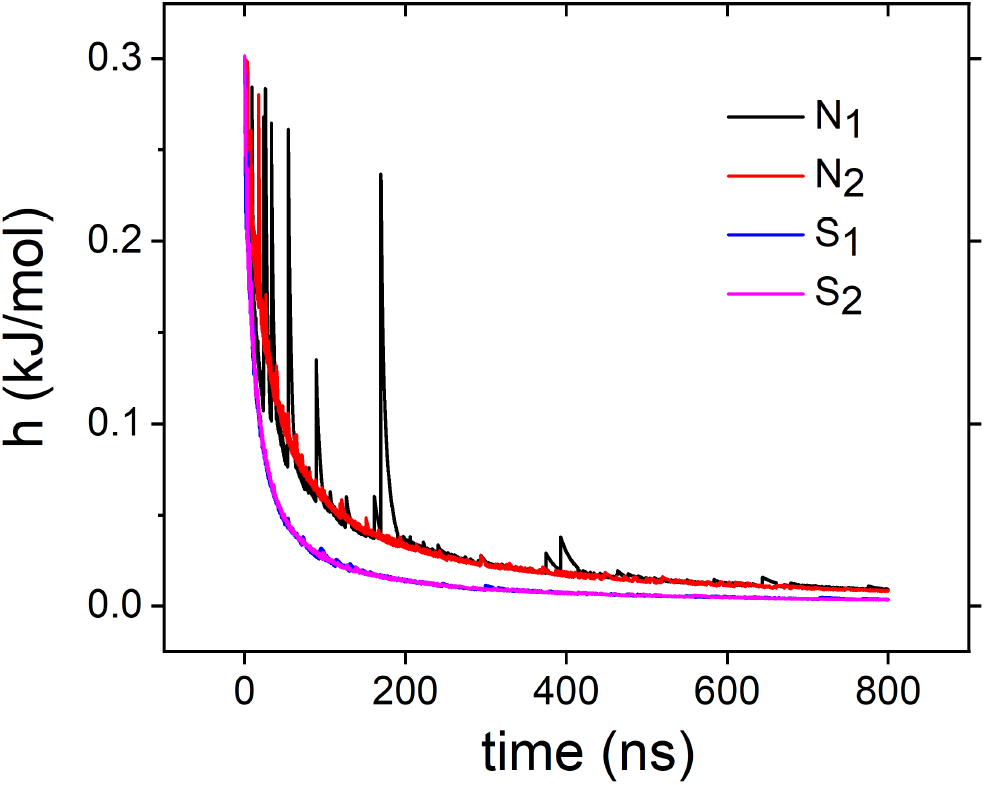
Height of the biasing Gaussian potentials (defined in Supplementary Equation (S2)) in the metadynamics as functions of simulation time. In our well-tempered metadynamics simulations, the initial value of the Gaussian height was set to 0.3kJ/mol, while the bias factor *η* = 200 was used to control the rate of potential decrease. After 200 ns, all the biasing Gaussians are smaller than 0.05kJ/mol, which further decrease to values less than 0.03kJ/mol after 300ns. These correspond to only about 1/10 of the initial values.

**Figure S9:**
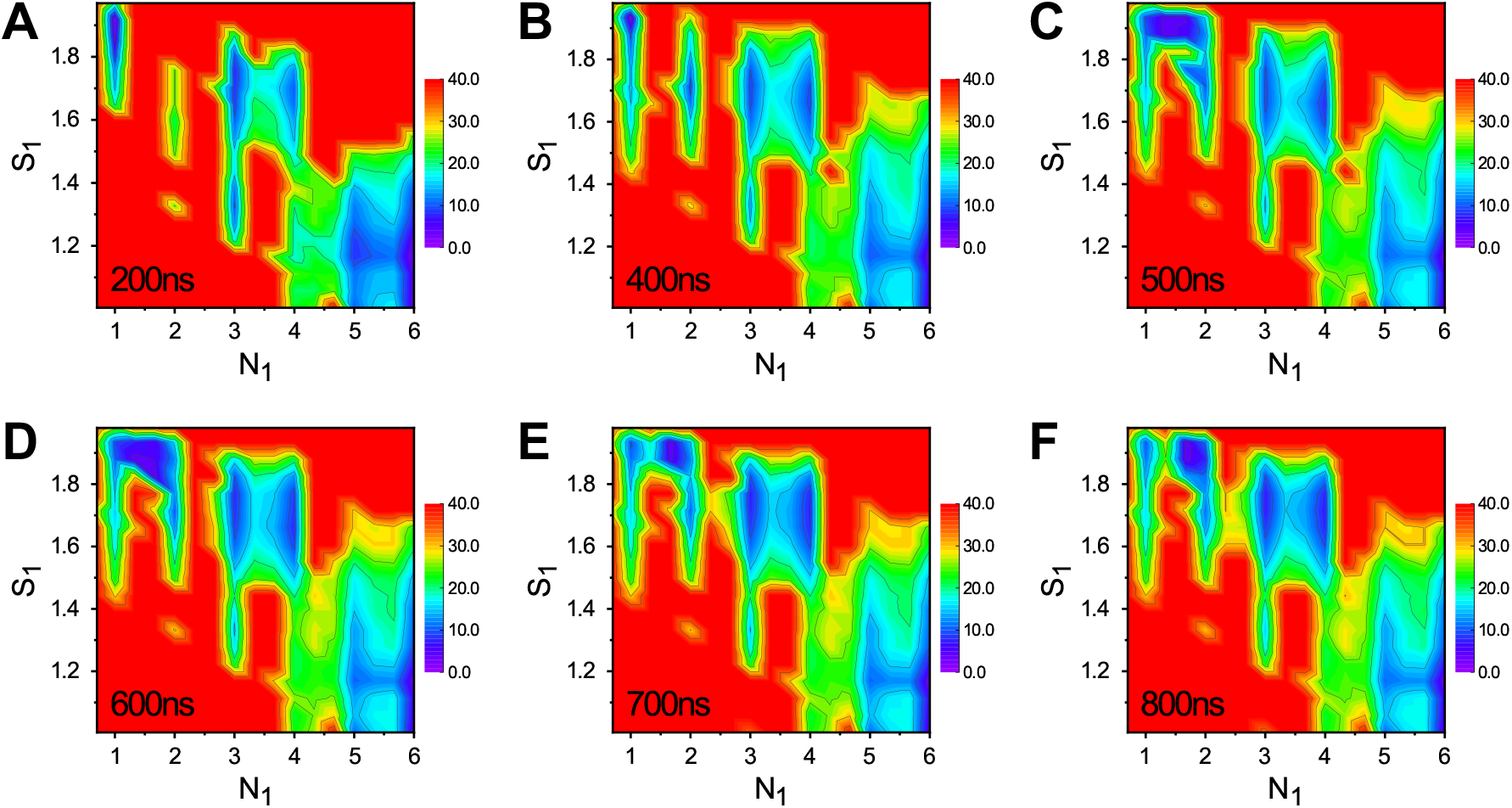
Time evolution of the two-dimensional *N*_1_-*S*_1_ free energy surface sampled by well-tempered bias-exchange metadynamics. Note that after 400 ns the two dimensional surfaces are stationary, indicating the convergence of the simulation results.

